# Apoptotic Cells May Drive Cell Death in Hair Follicles During their Regression Cycle

**DOI:** 10.1101/2020.11.16.384347

**Authors:** Bradley D. Keister, Kailin R. Mesa, Krastan B. Blagoev

## Abstract

Intravital microscopy in live mice have shown that the elimination of epithelial cells during hair follicle regression involves supra-basal cell differentiation and basal cell apoptosis through synergistic action of TGF-β (transforming growth factor) and mesenchymal-epithelial interactions. In this process the basal epithelial cells are not internally committed to death and the mesenchymal dermal papilla (DP) plays essential role in death induction. Because the DP cells are not necessary for completion of the cycle but only for its initiation it is still an open question what is the mechanism leading to the propagation of apoptosis towards the “immortal” stem cell population. Here, we use a quantitative analysis of the length of hair follicles during their regression cycle. The data is consistent with a propagation mechanism driven by apoptotic cells inducing apoptosis in their neighboring cells. The observation that the apoptosis slows down as the apoptotic front approaches the stem cells at the end of the follicle is consistent with a gradient of a pro-survival signal sent by these stem cells. An experiment that can falsify this mechanism is proposed.

## Introduction

Adult stem cells together with their supporting cells form stem cell niches which maintain the functionality of renewable tissue in different organs[1]. Stem cell niches have been identified in organs like the colon[2], the breast[3], the skin[4], the hair follicles[5], and the bone marrow[6]. Revealing the stem cell niche self-renewal dynamics is important not only for understanding tissue homeostasis, but also for understanding of initiation of cancer[7]. Furthermore the different stem-cell niche architecture in different organs may lead to a different rate of aging and susceptibility to cancer[8]. During the past decade significant advances were made in understanding stem cell niches thanks to the development of organoid cultures[9] and intravital microscopy[10].

Hair maintenance in mammals is governed by complex interactions between cells in and around the hair follicles[11]. Functional hair follicles cycle between growth (anagen, 3-5 years in humans), regression (catagen ~10 days in humans), and quiescence (telogen ~3 months in humans)[12]. At the beginning of telogen the hair falls off and after ~3 months new growth phase starts. In mice the phases are shorter, anagen lasts 1-3 weeks, catagen ~2 days, and telogen ~2 weeks[13]. During the regression phase most cells die, but a small set of stem cells don’t, and they replenish the follicle during growth. Recently[14] it was shown that extrinsic factors control cell apoptosis during that phase, but it is still an open question what is the origin of this signal. In addition, it is not understood why the apoptosis propagates along the follicle length rather than all cells dying simultaneously. The same study implicated the mesenchymal dermal papilla (DP) cells at the bottom of the follicle in the regression initiation through the release of a pro-apoptotic signal. This signal, associated with the transforming growth factor (TGF-β) can establish a spatial gradient along the hair follicle length inducing programmed cell death in the epithelial cell population. If such spatial signal is the sole cause of apoptosis it will lead to simultaneous apoptosis of all cells (with a rate proportional to the local concentration of TGF-β), which is in contradiction with the observations. During the regression phase, the DP cells follow the regressing cells until they reach the stem cell population in the hair follicle bulge. However, it was shown that while essential for initiating regression, the DP cells are not necessary for completion of the regression phase[14]. This suggests that only the DP signal released at the onset of the cycle is essential for regression and while the DP cells might continue releasing the signal until the end of the cycle it is not the only signal driving the apoptotic propagation. At the end of the cycle few stem cells remain viable in contact with the dermal papilla. This means that in addition to the apoptosis initiating DP signal(s), another mechanism is needed to explain the observed apoptotic propagation. In this paper we measured the length of hair follicles during catagen. The follicles were at different catagen stages and we obtained the follicle size distribution, which is consistent with a power law distribution. We also observed that, within a 12-hour period, shorter hair follicles regressed less than longer hair follicles, suggesting that the apoptotic propagation slows down as the dying cells approach the “immortal” stem cell pool. Here we propose a quantitative model according to which apoptotic cells release local signal priming their neighboring cells for apoptosis and the stem cells release a pro-survival signal setting a spatial gradient. This model is consistent with the experimentally measured distribution as well as with hair follicle regression deceleration.

## Results

### Short follicles regress slower than longer ones and basal epithelial cells don’t die until all cells before them are dead

A previous study[14] showed that the DP cells while needed for the initiation of the regression cycle are not necessary for its completion. To study the kinetics of follicle regression intravital microscopy was used in live mice to image 51 fluorescently labeled follicles and obtained the distribution of follicle lengths at two time points separated by 12 hours. The ordered-by-length set of imaged follicles is shown in Fig. 1a. In Fig. 1b the length change dependence on the initial length is shown. From this data it is clear that the regression rate of short follicles is less than the regression rate of longer follicles. During the regression cycle the death rate decreases as the follicles become shorter. During the regression cycle follicles regress independently and thus one can quantify the regression kinetics of a single follicle from the observation of an ensemble of follicles at a given time. Because the regressing follicles are independent and with variable length the follicles have started regressing at independent earlier times. A useful statistical tool for understanding the data is the probability distribution of follicle lengths. This distribution tells us what percent of follicles fall within a given range of lengths. To obtain this distribution we plot the percent of follicles within a given length range. The distributions at the two time points separated by 12 hours are shown in Fig. 2.

**Fig. 1.**
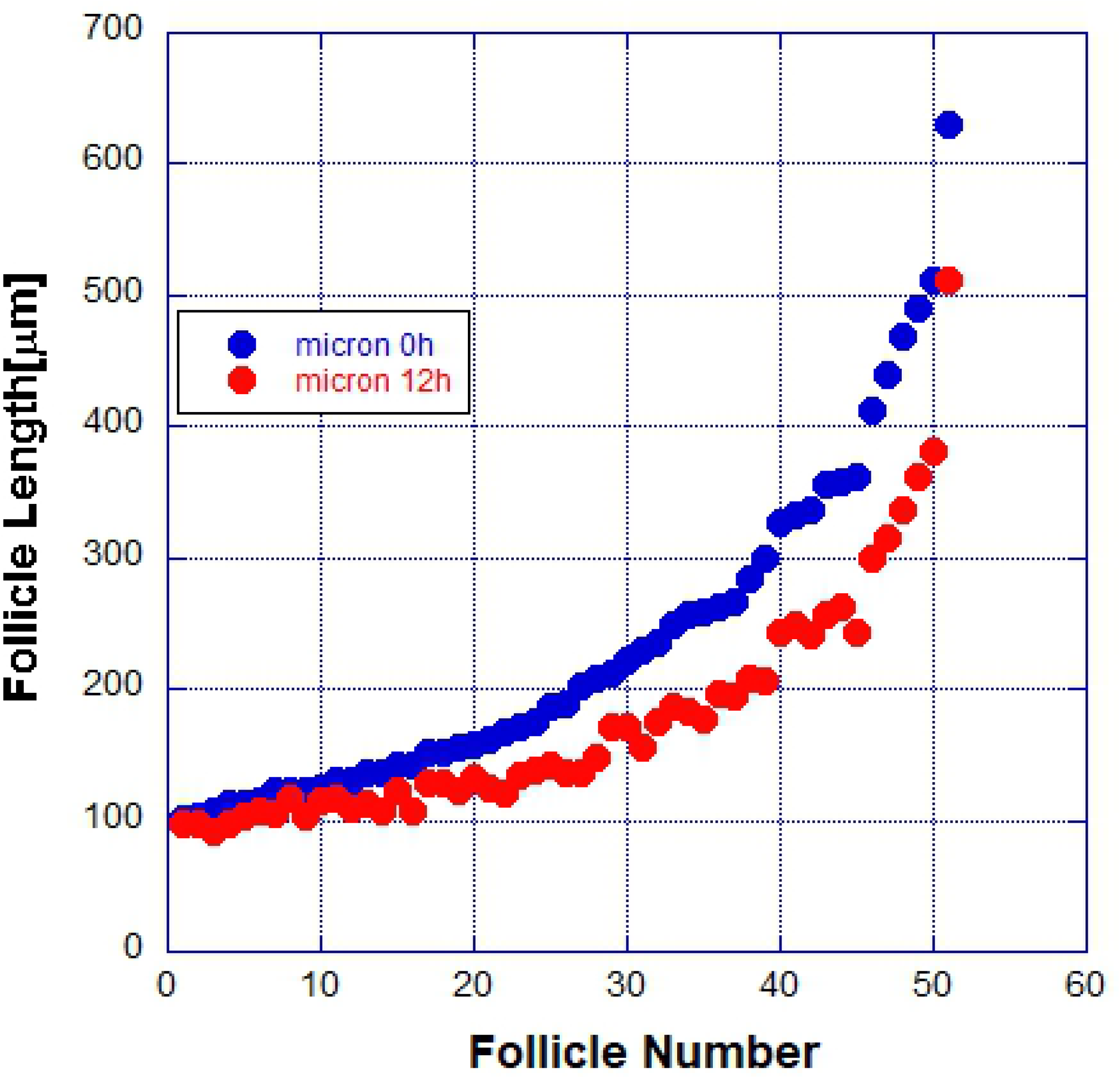
a) The length of 51 hair follicles was measured at two points of time separated by 12 hours. b) The dependence of the follicle length change on the initial length.

**Fig 2.**
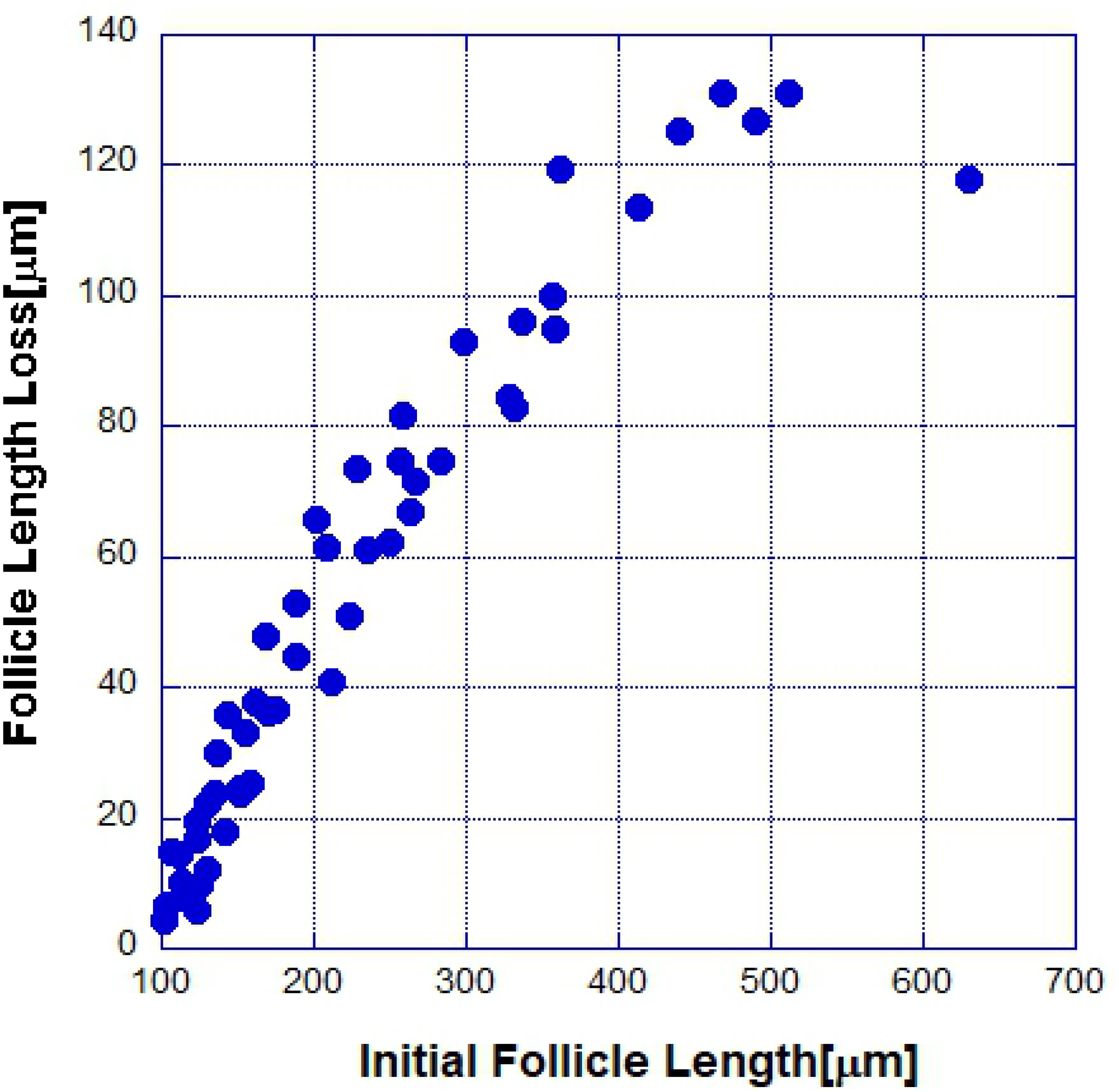
Percent of follicles with length within a small range (the probability distribution function).

We tried to fit these distributions to various simple functions and we found that the best fit was obtained with power law distribution (see Materials and Methods for a discussion of the fitting and possible problems). The observation that the DP cells are not needed for the completion of the regression cycle means that the initiation signal once released leads to the progressive cell death. Previous studies showed that the epithelial cells at a particular position along the follicle don’t die until all cells closer to the DP cells are dead[14].

### Apoptotic cell-cell communication can explain quantitatively the data

Here we introduce a mathematical model that can explain qualitatively the observed data. In this model the DP cells release an initiating signal, which primes the nearby epithelial cells for apoptosis. The dying cells release a local apoptotic factor which acts on the nearby cells and this cell death propagates further and further similar to the way fire propagates along a rope. In this model the apoptosis will propagate with a constant speed and it does not explain why the observed speed of follicle regression decreases and why the dying cells do not affect the stem cell pool at the end of the follicle. To model these observations we postulate that a pro-survival signal is released by the stem cell pool which degrades as the distance from the stem cell pool increases. To see if this model is quantitatively consistent with the observations, we modeled the described processes by a reaction-diffusion type equation[16]. We model the density, n(x,t), of dead cells along the length, x of the hair follicle at time t. The equation describing our model is:

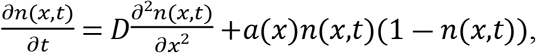

where x is the length along the follicle, t is time, and D is a diffusion coefficient describing the pro-apoptotic signal released by the dying cell. Here *a*(*x*) = 1 – exp (*k*(*x* – *L*)) describes a pro-survival factor released by the stem cell pool at the end (x = L) of the follicle and k is constant describing how fast along the follicle the pro-survival factor falls off. This model has only two fitting parameters: the diffusion coefficient D and the function a(x). Again, on general grounds this function is expected to decrease exponentially away from its source (the stem cell pool). In Fig. 3 we show the fit of this model to the data from Fig. 1a.

**Fig. 3.**
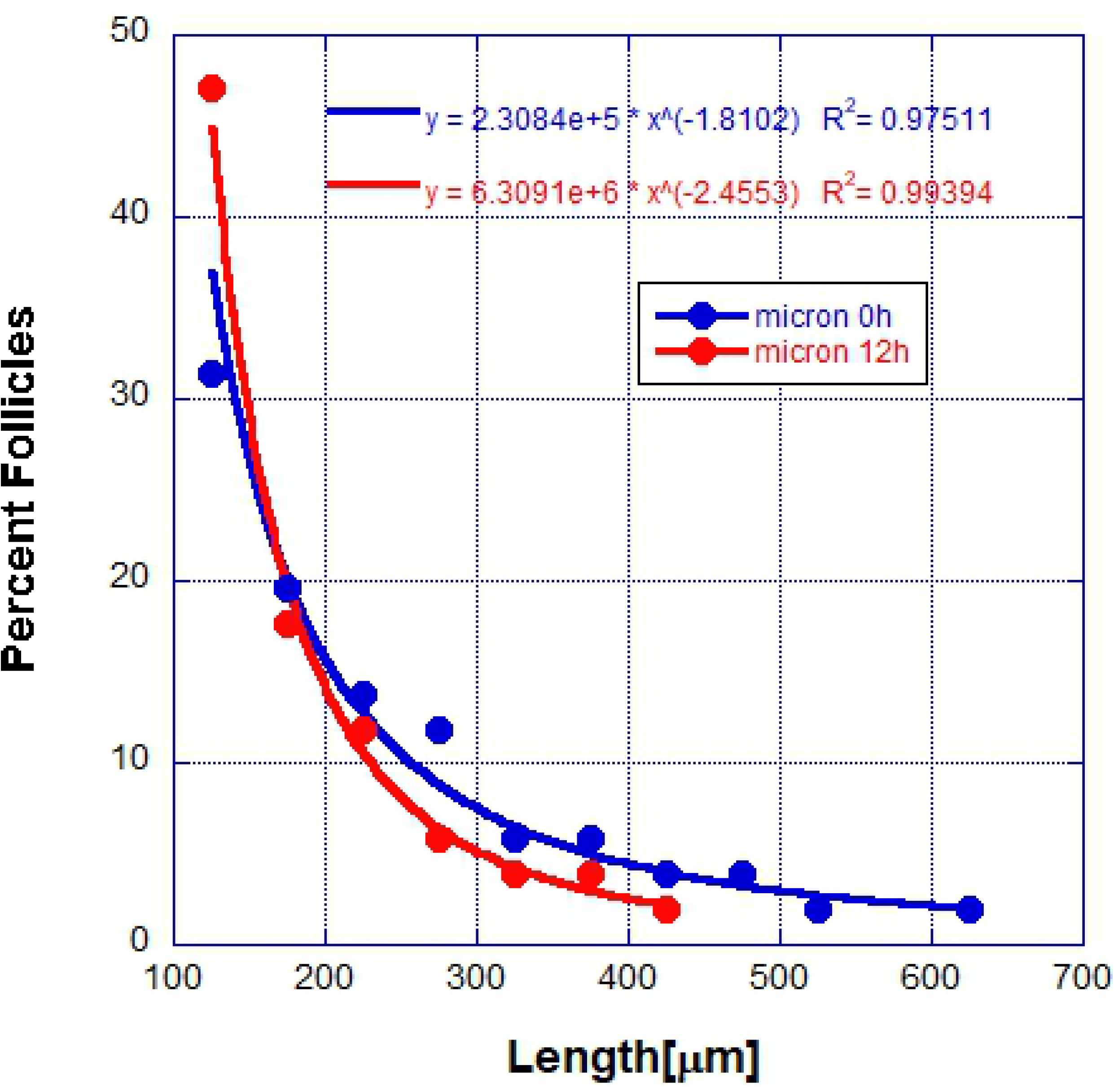
Simulations of the reaction-diffusion model (solid lines) and measured follicle length (solid circles).

Because of the putative pro-survival factor possibly released by the stem cells, the propagation of apoptosis along the follicle becomes slower and slower as one approaches the stem cells and it stops when it reaches them. In Fig. 4 the propagation of the initial cell death along the follicle is shown.

**Fig. 4.**
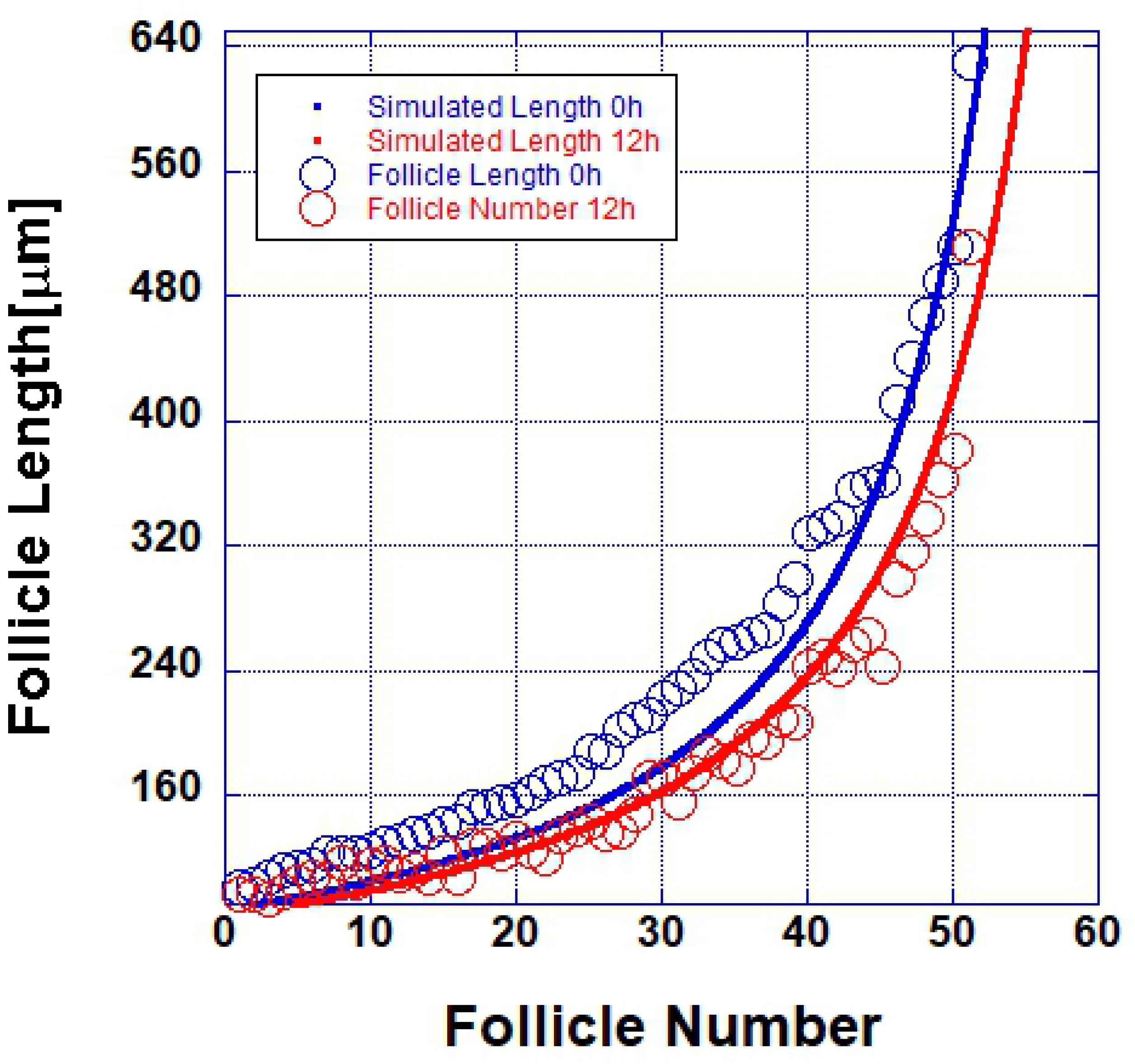
Cell extinction slows down as the stem cells at the end of the follicle are approached in the mathematical model as observed in the set of measured hair follicles.

The proposed model for the follicle regression cycle produces a power law distribution of follicle lengths. While the experimental data suggests a power law distribution, the follicles are within their biological lengths and mathematically it is impossible to have high confidence that the distribution is a power law. In the case of the model we have many more data points and we are confident in the power law distribution. We observed a typical power law distribution generated by our model (Fig. 5)

**Fig. 5.**
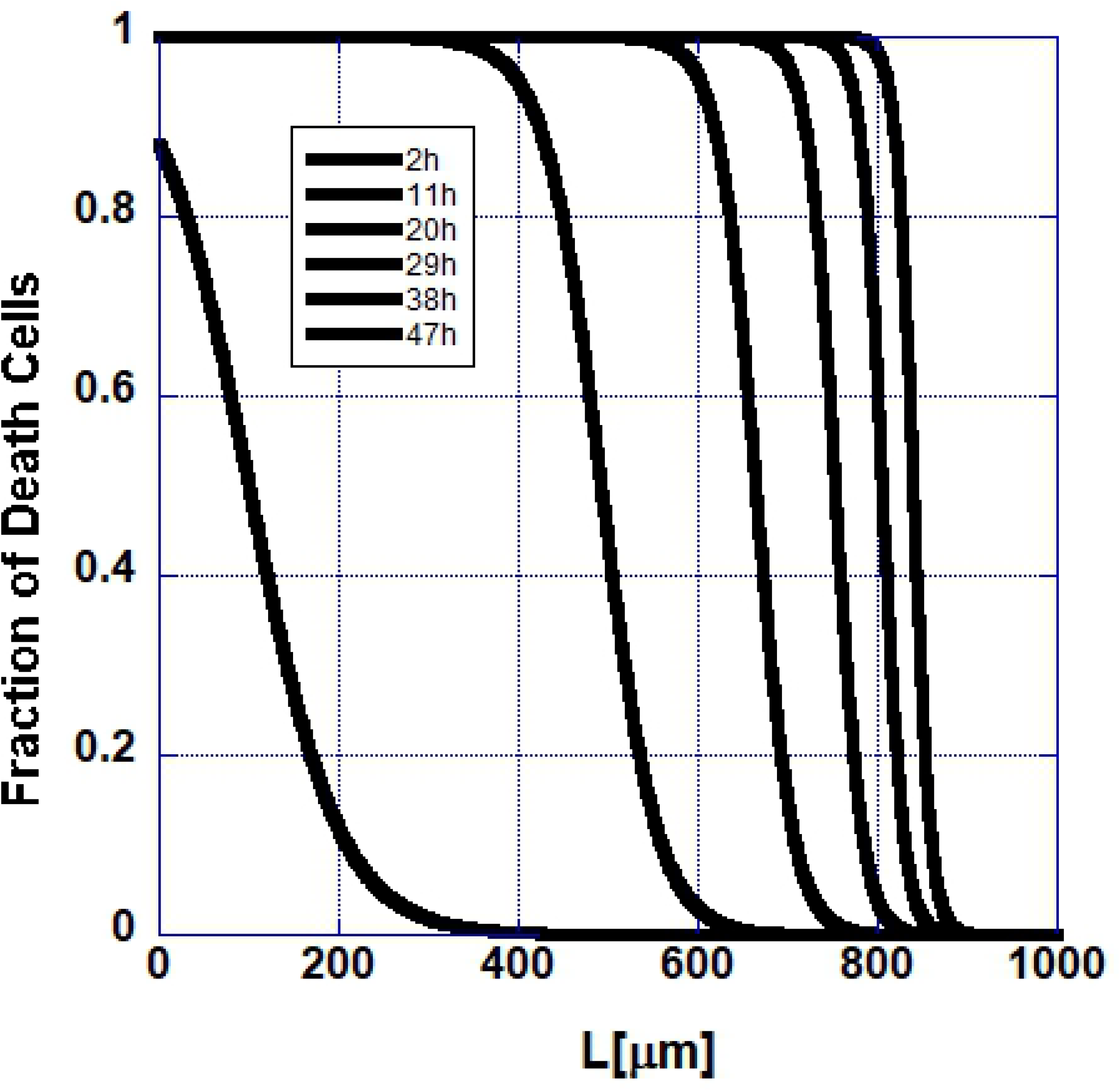
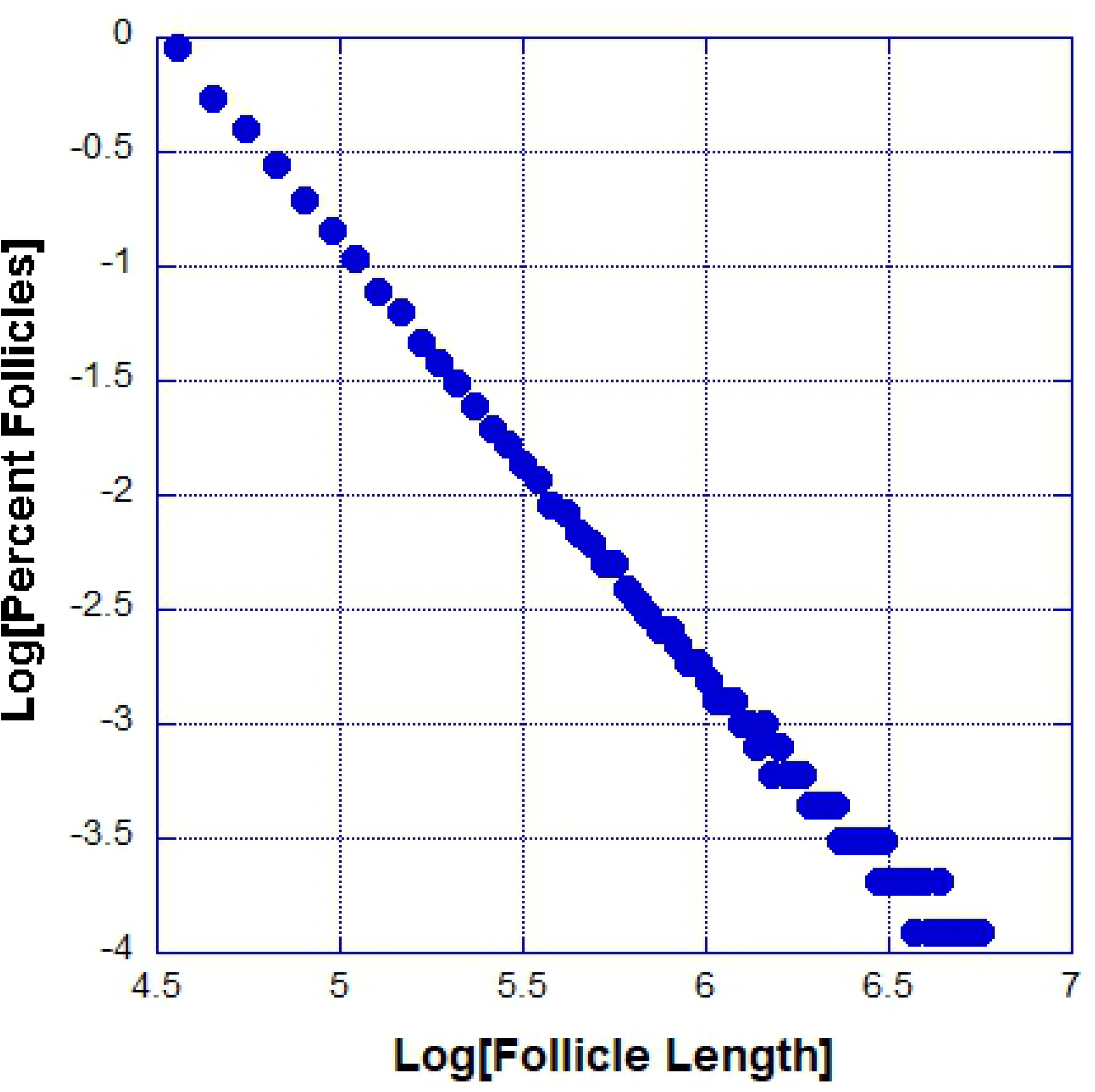
The power law probability distribution function of follicle sizes generated by our model.

So far, we have shown that a simple deterministic model of cell-cell communications is consistent with the observations and that a model in which the apoptosis is induced by a spatially separated cellular compartment is not. The next question we asked was how sensitive to noise our results are. We introduced multiplicative noise in the reaction term drawn from a uniform distribution. The results were not sensitive to such noise and the dynamics was self-averaging.

## Discussion

In this paper we introduced a mathematical model of the hair follicle regression cycle that postulates that the regression is initiated by the dermal papilla, but that this signal affects only the cells adjacent to it. The subsequent regression propagates due to the death of cells, which induce apoptosis in their neighbors. This model is consistent with the observation that the dermal papilla is not necessary for the completion of the regression cycle. The mathematical model is based on a reaction-diffusion-type dynamics in which the reaction term strength, describing the cell-cell induced apoptosis, decreases towards the stem cells most likely due to the release by the stem cells of pro-survival diffusible factor, slowing down the effect of dying cells on their neighbors. In the regression mechanism proposed here apoptotic cells induce apoptosis in their neighbors and this signal propagates forward along the hair follicle and our prediction is that ablation of cells ahead of the apoptotic propagation front will halt the follicle regression, while in a model where external diffusing signal causes the regression ablation is expected to have little effect on the regression cycle. Such experiments can falsify the mechanism proposed here.

In conclusion, hair follicle regression may be governed by cell-cell induced programmed cell death, which slows down as the stem cell compartment is approach and does not affect the stem cell compartment from which the growth phase is initiated.

## Materials and Methods

### Experimental procedures and mice models

The intravital microscopy and all the experimental methods, procedures and mice have been described in detail elsewhere[14].

### Mathematical Modeling

The reaction-diffusion equation describing our model is an initial-value problem solved via Euler’s method for the time dependence, using a spatial discretization of n(x,t). We use two approaches to check the numerical calculations. The first is to replace Euler’s method with fourth-order Runge-Kutta integration, and/or to compare the use of fourth-order vs second-order evaluation of the Laplacian on the lattice. For sufficiently short time steps, the results from these methods are practically identical. The second check uses the fact that, when the signal a(x) is set to a constant, the solution becomes a time-independent shape that propagates with a velocity determined by the other parameters. We wrote a separate code to produce such shapes, and confirmed that, when used as an initial condition for the code with damping, the shape propagates without change.

## ACKNOWLEDGMENTS

We would like to thank Valentina Greco for valuable discussions. The National Science Foundation supported the work of K. B. Blagoev and Bradley Keister through the NSF IR/D program. The National Science Foundation had no role in the study design, data collection and analysis, decision to publish, or preparation of the manuscript. The views presented here are not those of the National Science Foundation and represent solely the views of the authors.

